# Neural prediction decorrelation reveals that adversarial robustness substantially improves DNN prediction accuracy across the entire human auditory cortex

**DOI:** 10.64898/2026.08.05.743059

**Authors:** David Skrill, Jenelle Feather, Sam V. Norman-Haignere

## Abstract

Sensory neuroscientists seek to model the neural computations that encode complex stimuli. Distinct encoding models often make similar predictions for natural stimuli such as speech, posing a challenge for model comparison. We developed a method to synthesize stimuli that decorrelate model predictions across a neural population, termed neural prediction decorrelation (NPD). Using fMRI responses to NPD sounds, we compared standard and adversarially robust deep neural network models of human auditory cortex. Prediction accuracy for NPD sounds was substantially better for the adversarially robust model in every region tested, an effect completely masked with natural sounds. Population responses to natural and synthesized NPD sounds shared an interpretable low-dimensional organization that was reproduced by the robust encoding model. NPD provides a general approach for comparing encoding models and reveals that adversarial robustness expands the predictive power of DNNs beyond natural stimuli, which is likely critical for targeting population activity through stimulus synthesis.

## Introduction

A central goal of sensory neuroscience is to model how neural populations encode complex stimuli (Pouget et al. 2000; Wu et al. 2006; de Heer et al. 2017; Wang et al. 2025). Encoding models are often evaluated and compared by measuring their ability to predict neural responses to natural stimuli (Naselaris et al. 2011; Yamins et al. 2014; Schrimpf et al. 2020), such as speech (de Heer et al. 2017; Kell et al. 2018; Brodbeck et al. 2018; Hamilton et al. 2021; Keshishian et al. 2023). A key challenge is that distinct models often show similar neural prediction accuracy (Norman-Haignere and McDermott 2018; Groen et al. 2018; Tuckute et al. 2023; Canatar et al. 2023; Conwell et al. 2024). This similarity is often attributed to shared structure across models (Huh et al. 2024; Storrs et al. 2021) or limitations of the neural measurements (e.g., coarse neuroimaging data) (Kanwisher 2025; Norman-Haignere et al. 2022; O’Connell and Kelly 2021), but it can also arise from the stimulus set itself. If models are only tested on stimuli that elicit similar model predictions, then neural data cannot reveal which model is better (Norman-Haignere and McDermott 2018; Groen et al. 2018). This problem is pervasive when modeling neural responses to complex stimuli (DiLiberto et al. 2015; Norman-Haignere and McDermott 2018; Groen et al. 2018; Daube et al. 2019; Tuckute et al. 2023; Conwell et al. 2024), and developing effective solutions would have broad significance.

Deep neural networks (DNNs) trained on challenging tasks have been found to predict neural responses better than traditional feature sets across multiple sensory and cognitive systems (Kell et al. 2018; Tuckute et al. 2023; Li et al. 2023; Rupp et al. 2025; Eickenberg et al. 2017; Güçlü and van Gerven 2015; Yamins et al. 2014; Wang et al. 2025; Caucheteux and King 2022; Schrimpf et al. 2021; Goldstein et al. 2022; Jain and Huth 2018). Because modern deep learning has produced many models with different architectures and training procedures, comparing their neural prediction accuracy has become an important way to test computational hypotheses (Kell and McDermott 2019; Saxe et al. 2021) and assess which models best align with the brain (Kietzmann et al. 2019; Schrimpf et al. 2020; Geirhos et al. 2018; Feather et al. 2023). A current challenge, however, is that many DNNs predict neural responses with similarly high accuracy (Schrimpf et al. 2020; Storrs et al. 2021; Tuckute et al. 2023; Conwell et al. 2024), even for networks whose behavior diverges from biological systems. For example, DNNs often make confident recognition judgments for stimuli that are unrecognizable to human observers (Feather et al. 2023), and small perturbations to a natural stimulus (“adversarial attacks”) can dramatically alter a network’s decisions without altering human perception (Goodfellow et al. 2015; Carlini and Wagner 2018). Training DNNs to be robust to such adversarial attacks improves the alignment between human and model decisions (Feather et al. 2023; Gaziv et al. 2023; Engstrom et al. 2019). Yet, similar to prior work (Feather et al. 2023), we find that standard and robust DNNs predict human cortical responses to natural sounds with similar accuracy.

We hypothesized that this similarity in neural prediction accuracy reflects a limitation of the stimulus set. Consistent with this hypothesis, we find that robust and standard models make virtually indistinguishable predictions for natural sounds. To address this problem, we developed a method to synthesize stimuli that decorrelate model predictions across a target neural population, which we term neural prediction decorrelation (NPD). Unlike prior behavioral methods (Wang and Simoncelli 2008; Golan et al. 2020), NPD is directly applicable to high-dimensional neural responses and can be used with any sensory modality (e.g., audition, vision) or recording method (e.g., functional MRI, electrophysiology).

We applied NPD to compare standard and adversarially robust models of the human auditory cortex using functional MRI (fMRI). NPD synthesized a new sound set that strongly decorrelated predictions across the auditory cortex, including for held-out participants. We then measured fMRI responses to these NPD sounds in new participants and found that the robust model retained strong prediction accuracy, whereas the standard model’s prediction accuracy fell to near zero, unmasking a large performance difference in every tested region. Population responses to natural and NPD sounds were organized in a shared low-dimensional subspace with interpretable structure. Robust models fit using only natural sounds were able to predict this shared population structure, demonstrating strong generalization beyond natural stimuli.

## Results

### Encoding models derived from standard and robust DNNs predict human cortical responses to natural sounds with similar accuracy

We first compared the neural prediction accuracy of auditory fMRI encoding models derived from pretrained standard and adversarially robust DNNs. We tested a standard DNN that previously showed good prediction accuracy of fMRI responses to natural sounds (CochResNet50, Feather et al. (2023)). This model consists of a time-frequency cochleagram front-end (Glasberg and Moore 1990; McDermott and Simoncelli 2011) followed by a convolutional architecture with residual connections (ResNet50 (He et al. 2016)), and was trained to recognize words in the presence of complex background sounds. Prior work found that this model predicts fMRI responses to both speech and non-speech natural sounds better than standard acoustic models and replicates hierarchical functional organization (Tuckute et al. 2023). We compared this standard DNN to an adversarially robust DNN that was identical in its architecture and training, but was additionally trained to be robust to adversarial perturbations applied to the input cochleagram representation (*l*^2^-norm constraints: *ϵ* = 1.0, (Madry et al. 2018; Feather et al. 2023)).

We used activations from the pretrained DNNs to predict human fMRI responses to 165 natural sounds (each 2 seconds) (Fig. 1A) collected from 30 different participants across two prior studies (Norman-Haignere et al. 2015; Boebinger et al. 2021), henceforth referred to as NH2015 and B2021, respectively. Following prior studies (Norman-Haignere et al. 2015; Boebinger et al. 2021), we measured the time-averaged response of each voxel to each of the 165 sounds, since the temporal resolution of fMRI is coarse relative to the sound’s duration. We then predicted the response of each voxel as a weighted sum of DNN activations computed for the same 165 natural sounds. As has become standard, we used cross-validated ridge regression to compute the regression weights and select the DNN layer that best predicted each voxel’s response (Naselaris et al. 2011; Güçlü and van Gerven 2015; Kell et al. 2018; Santoro et al. 2014; de Heer et al. 2017; Millet et al. 2022; Tuckute et al. 2023). We measured neural prediction accuracy as the Pearson correlation between the measured and model-predicted response for each voxel, separately for each model (Fig. 1B, left).

**Fig. 1.**
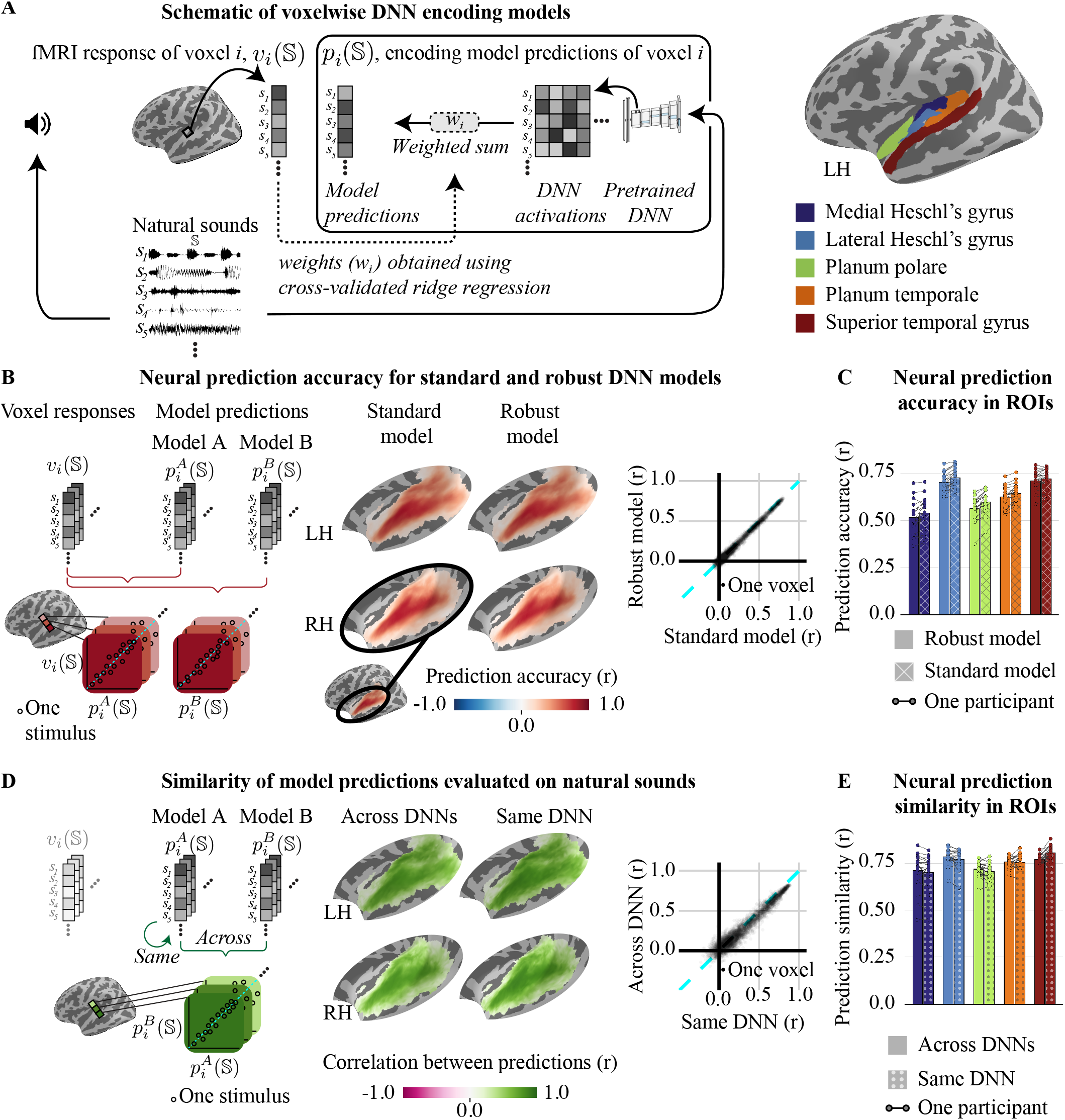
Similar fMRI prediction accuracy for standard and robust DNN models across human auditory cortex. **A** We measured time-averaged fMRI responses to a set of natural sounds (S), producing a vector of responses across the sound set for each voxel (*v*_*i*_(S)). DNN activations were then measured for this same sound set using a pretrained DNN model. Voxel responses were predicted as a weighted sum of DNN activations from the best predicting DNN layer for each voxel. The weights were inferred using cross-validated and regularized regression. This procedure yields a vector of model predictions across the sound set for each fMRI voxel in the auditory cortex (*p*_*i*_(S)). **B** Voxelwise fMRI prediction accuracy was measured for standard and adversarially robust DNN models by correlating the measured and model-predicted response of each voxel. This figure plots the correlation for each voxel averaged across participants in standardized coordinates (FsAverage) (data from individual participants is plotted in Supplemental Fig. S1). The scatter to the right directly compares the prediction accuracy of robust (y-axis) vs. standard (x-axis) models for each voxel (one dot: one voxel). **C** Neural prediction accuracy for robust (solid) and standard (hashed) models across different anatomical regions-of-interest (ROIs). Each circle plots the median prediction accuracy across all voxels in each ROI from both hemispheres in each participant. Lines connect prediction accuracies for standard and robust models from the same participant. Bars plot the average across participants. **D** Similarity of standard and robust model predictions for the natural sounds tested in the experiment. Similarity was measured by correlating the model predictions across models for each voxel (“Across DNNs”, left). The robust and standard encoding models were fit to fMRI responses using two, non-overlapping sets of natural sounds. This allows us to plot a measure of the maximum possible correlation, given the fit to noisy neural data, by correlating the predictions from the same DNN (standard vs. standard and robust vs. robust) fit using two different stimulus sets (“Same DNN”, right). The scatter to the right directly compares the Across DNN correlation (y-axis) vs. the Same DNN correlation (x-axis) for each voxel (one dot: one voxel). **E** Similarity of model predictions across DNN models (solid bars) and within the same model (dotted bars) for different anatomical ROIs. The format is otherwise the same as panel C.

We found that the neural prediction accuracy for the standard and robust models was very similar (Fig. 1B; see Fig. S1A for individual participant maps) across all anatomical regions of the auditory cortex (Fig. 1C). We used a simple metric 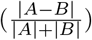 that ranged from 0 (identical) to 1 (maximally different) to quantify the dissimilarity of neural prediction accuracies *A* and *B* (e.g., of the standard and robust models), and bootstrapped this metric across participants to estimate 95% confidence intervals. Dissimilarity metrics were close to 0 for all regions of the auditory cortex (Medial Heschl’s gyrus: 0.023 (0.013, 0.033) Lateral Heschl’s gyrus: 0.017 (0.011, 0.022) Planum polare: 0.031 (0.021, 0.041) Planum temporale: 0.015 (0.006, 0.023) Superior temporal gyrus: 0.007 (0.001, 0.013)). Thus, even though prior behavioral evidence suggests that adversarial robustness improves the alignment between models and human decisions (Tsipras et al. 2019; Feather et al. 2023; Gaziv et al. 2023), adversarial robustness does not substantially alter neural prediction accuracy when evaluated using natural sounds.

### NPD sounds decorrelate model predictions across the entire human auditory cortex

We next investigated whether similar neural prediction accuracy between standard and robust models could be explained by similar model predictions for the natural sounds tested in the experiment. We measured similarity by correlating the model-predicted fMRI response to natural sounds between robust and standard encoding models (Fig. 1D). We compared this “Across DNN” correlation with a “Same DNN” correlation that estimates the similarity one would expect when comparing predictions from the same pretrained DNN, but fit using noisy neural responses. Specifically, we first fit fMRI encoding models from both standard and robust DNNs using two non-overlapping sets of natural sounds. We then correlated the predictions for a third non-overlapping sound set, either across DNN models (robust-standard) or within the same DNN model (robust-robust, standard-standard) (see Methods for details). We found that the Across DNN correlation was very similar to the Same DNN correlation (Fig. 1D; see Fig. S1B for individual participant data) in all anatomical regions of the auditory cortex (Fig. 1F) with dissimilarity metrics close to 0 (Medial Heschl’s gyrus: 0.006 (0.000, 0.017) Lateral Heschl’s gyrus: 0.009 (0.003, 0.015) Planum polare: 0.009 (0.003, 0.016) Planum temporale: 0.001 (0.000, 0.011) Superior temporal gyrus:0.022 (0.013, 0.029)). This finding indicates that the predictions from standard and robust models are about as similar as would be expected when comparing two identical models.

These results raise the possibility that similar neural prediction accuracy reflects a limitation of the experimental stimuli. Specifically, we wondered whether it would be possible to design a targeted stimulus set for which the models make distinct predictions, and if so, whether one model would show higher neural prediction accuracy for those stimuli. To answer this question, we developed an approach to synthesize stimuli for which competing models make reliably distinct predictions for all responses in a neural population (Fig. 2A). Nothing about our method is specific to DNNs, sounds, or fMRI, and the method can be applied to more than two competing models. We therefore describe the method in general terms, using our application of comparing standard and robust fMRI encoding models as a concrete example. We use the word “channel” to refer to individual measurement sites in a neural population (e.g., fMRI voxels).

**Fig. 2.**
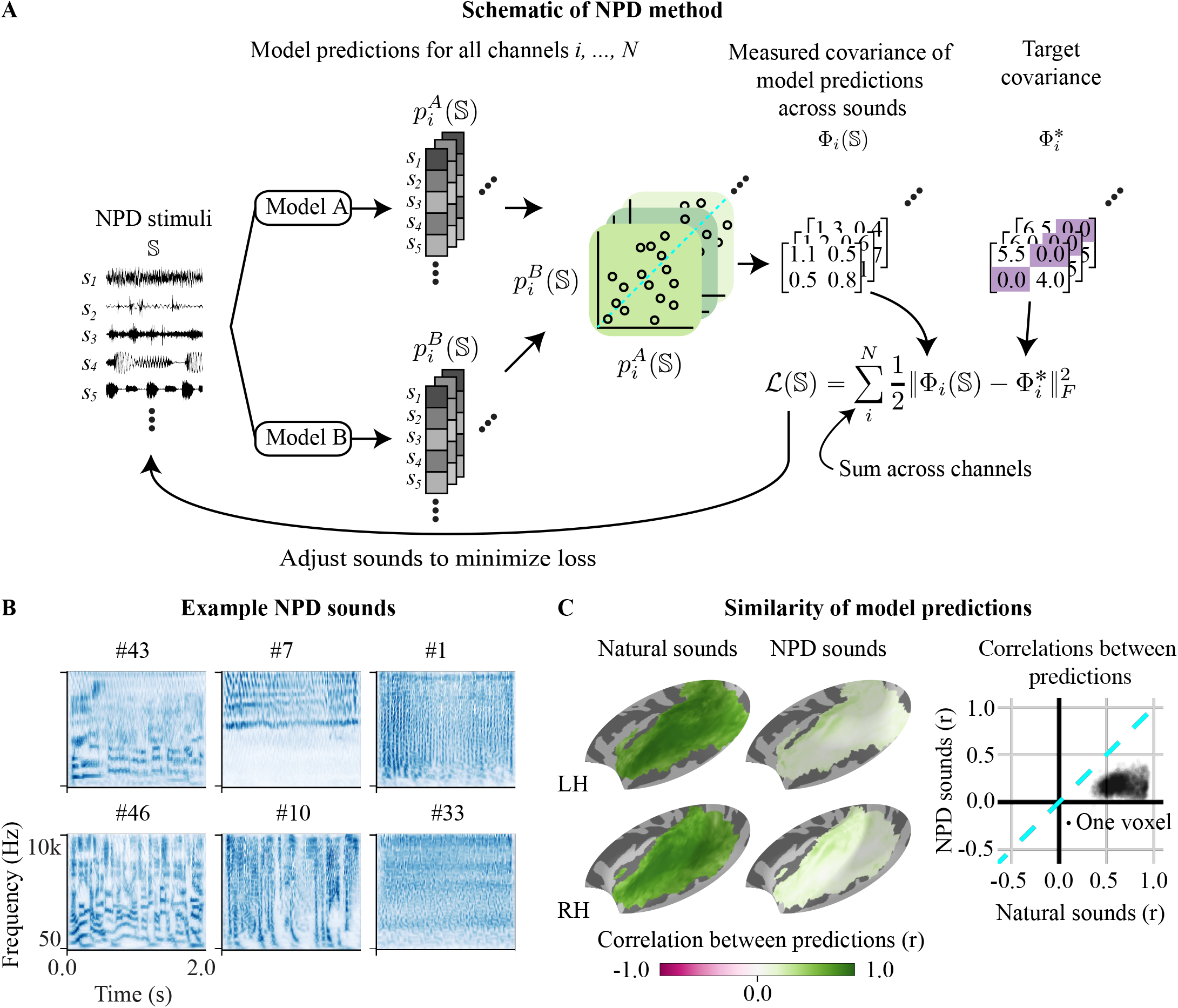
Neural prediction decorrelation. **A** Schematic of the procedure used to synthesize NPD stimuli for the special case of two models (A and B, e.g., robust and standard models). The algorithm computes encoding model predictions 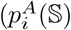 and 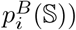 for every channel (*i*) in a neural population (here fMRI voxels) for a set of synthesized stimuli (S). These predictions are used to compute a [model × model] covariance matrix (Equation 1) for each channel. This covariance matrix measures the variance/covariance of the model predictions across the synthesized stimulus set. The synthesis algorithm drives this empirically measured covariance matrix to match a target covariance matrix (Equation 4) by minimizing a loss function that measures the distance (squared Frobenius norm) between the measured and target covariance matrix for each channel, and then summing these distances across channels. The off-diagonal elements of the target covariance matrix (highlighted in purple) are set to 0, which drives the correlation of the model predictions to 0. The on-diagonal elements are set to a high value to preserve high response variance (here, 5× the model-predicted variance for natural stimuli). Gradients from the loss function are backpropagated to the stimulus and used to update the stimulus set to minimize the loss. **B** Time-frequency cochleagrams for example NPD sounds. Cochleagrams plot energy as a function of time and frequency, similar to a spectrogram, but computed using filters and envelope compression designed to coarsely mimic that present in the peripheral auditory system. The sound set was synthesized to collectively decorrelate fMRI voxel predictions between standard and robust DNN models across the human auditory cortex. Sounds can be listened to at https://labsites.rochester.edu/neural-prediction-decorrelation/. **C** Similarity of model predictions for natural vs. NPD sounds. Correlations between standard and robust DNN model predictions, computed separately for natural (left map) and NPD sounds (right map). The scatter plot compares correlations for NPD (y-axis) vs. natural (x-axis) sounds for each voxel in the auditory cortex (data averaged across participants; individual participant data are plotted in Supplemental Fig. S2). Notably, the participants used to synthesize the NPD sounds were excluded from this analysis. Thus, this analysis tests whether our synthesis procedure decorrelates predictions in new subjects never used to synthesize the sounds.

In our framework, each encoding model (*m* ∈ *M* ) defines a mapping between any stimulus (*s*_*j*_) and the model-predicted response 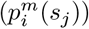 to that stimulus for each channel (*i*) in the neural population. Our goal is to synthesize a new stimulus set (S) that decorrelates predictions between models. Thus, we optimize the stimulus set given a predefined set of encoding models, in contrast with the more common setting in which models are optimized given fixed stimuli.

Our optimization objective decorrelates model predictions by driving the covariance between the predictions of different models towards zero. At the same time, the optimization algorithm encourages predictions from all models to maintain high variance, which is useful because low stimulus-driven variance reduces signal-to-noise in neural data. The variances and covariances of the model predictions are captured by a covariance matrix (Equation 1; 2 × 2 for the case of two models), whose diagonal and off-diagonal elements reflect the variances (Equation 2) and covariances (Equation 3), respectively, for the stimulus set:

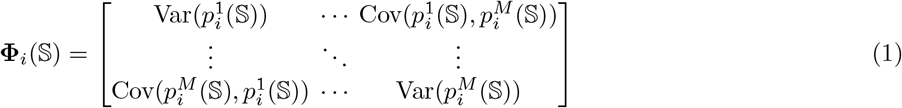

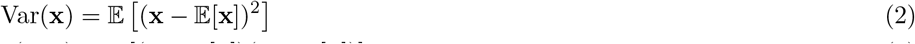

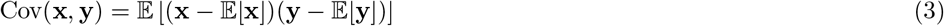

The optimization algorithm finds a stimulus set with a desired covariance structure for each channel in the population. Specifically, for each channel, we measure the distance between the empirical covariance matrix (**Φ**_*i*_(S)) and a target covariance matrix 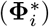:

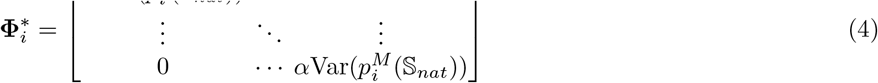

The off-diagonal elements of this target covariance matrix are set to 0, thus driving the covariance, and therefore the correlation, of the empirical model predictions to 0. The diagonal elements are set to a high value to preserve high response variance. In our study, we set these target variances to 5 times the model-predicted variance for natural sounds 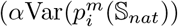, where *α* = 5). The goal was not to achieve this exact variance value, but rather to ensure high variance in the neural and predicted responses to synthesized sounds, which we confirmed empirically.

The complete loss function sums these distances across all channels in the population, encouraging the empirical covariance matrix to match the target covariance matrix for all channels in the population:

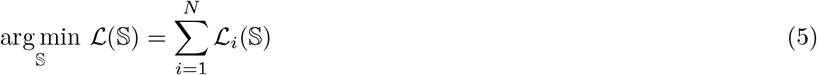

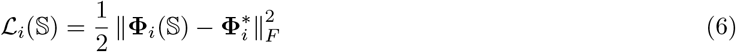

We refer to the stimuli generated through this optimization procedure as “Neural Prediction Decorrelation” (NPD) stimuli, as they are designed to simultaneously decorrelate the model predictions for all channels in a target neural population.

We used this approach to synthesize 60 NPD sounds that decorrelated model predictions between robust and standard encoding models across all voxels in the human auditory cortex. We initialized these 60 NPD sounds using unstructured Gaussian noise and performed gradient descent on the sounds to minimize equation 5. This optimization produced a diverse set of complex sounds that were acoustically and perceptually distinct from each other despite being initialized as unstructured noise (Fig. 2B; sounds can be listened to at the webpage supporting the work: https://labsites.rochester.edu/neural-prediction-decorrelation). Some sounds had a speech-like quality, although they were clearly distinct from natural speech without intelligible words or semantic content (e.g., NPD sounds #46 and #43 in Fig. 2B). Other sounds had tone-like (e.g., #7) or noise-like qualities (e.g., #33), and many exhibited complex patterns of temporal (e.g., #1) or spectrotemporal modulation (e.g., #10).

We next tested whether the NPD sounds decorrelated model predictions in held-out participants from a separate study. We synthesized NPD sounds using voxelwise encoding models from participants in one study (NH2015; 10 participants)(Norman-Haignere et al. 2015), and then evaluated the sounds using non-overlapping participants from a second study (B2021; 20 participants)(Boebinger et al. 2021). For the B2021 participants, we fit standard and robust encoding models using voxel responses to natural sounds. Using the fit models, we computed predictions for both NPD sounds and an independent set of natural sounds that were not used for fitting. We then correlated the predictions between robust and standard encoding models, separately for the NPD and natural sounds. Across auditory cortex, the correlation dropped substantially both at the group level (Fig. 2C) and within individual participants (Fig. S2A). The group-level median correlation across voxels was approximately eightfold lower for NPD sounds (0.102) than for natural sounds (0.797) (Fig. 2C; see S2). At the group level, every voxel in the auditory cortex with a reliable response to natural sounds showed a lower correlation for NPD than for natural sounds, even though the participants used for evaluation were entirely separate from those used for synthesis. Model-predicted variances were also high for the NPD sounds relative to natural sounds, as desired (Fig. S5).

### NPD sounds reveal that adversarial robustness substantially improves neural prediction accuracy

We conducted a new experiment (study S2026) in non-overlapping participants in which we measured fMRI responses to the 60 synthesized NPD sounds as well as 120 natural sounds (Fig. 3) (N=9 participants; the 120 natural sounds were a subset of the 165 natural sounds used in the NH2015 and B2021 studies). Overall, voxel responses to the NPD sounds were as strong and varied as those to natural sounds throughout primary and non-primary auditory cortex (Fig. S4). To test whether the NPD sounds were useful for model comparison, we fit robust and standard voxel-wise encoding models in these new participants using 60 natural sounds. We then evaluated neural prediction accuracy on the 60 NPD sounds and the remaining 60 held-out natural sounds by correlating the measured and model-predicted responses, separately for each model (standard and robust) and sound set (NPD and natural).

**Fig. 3.**
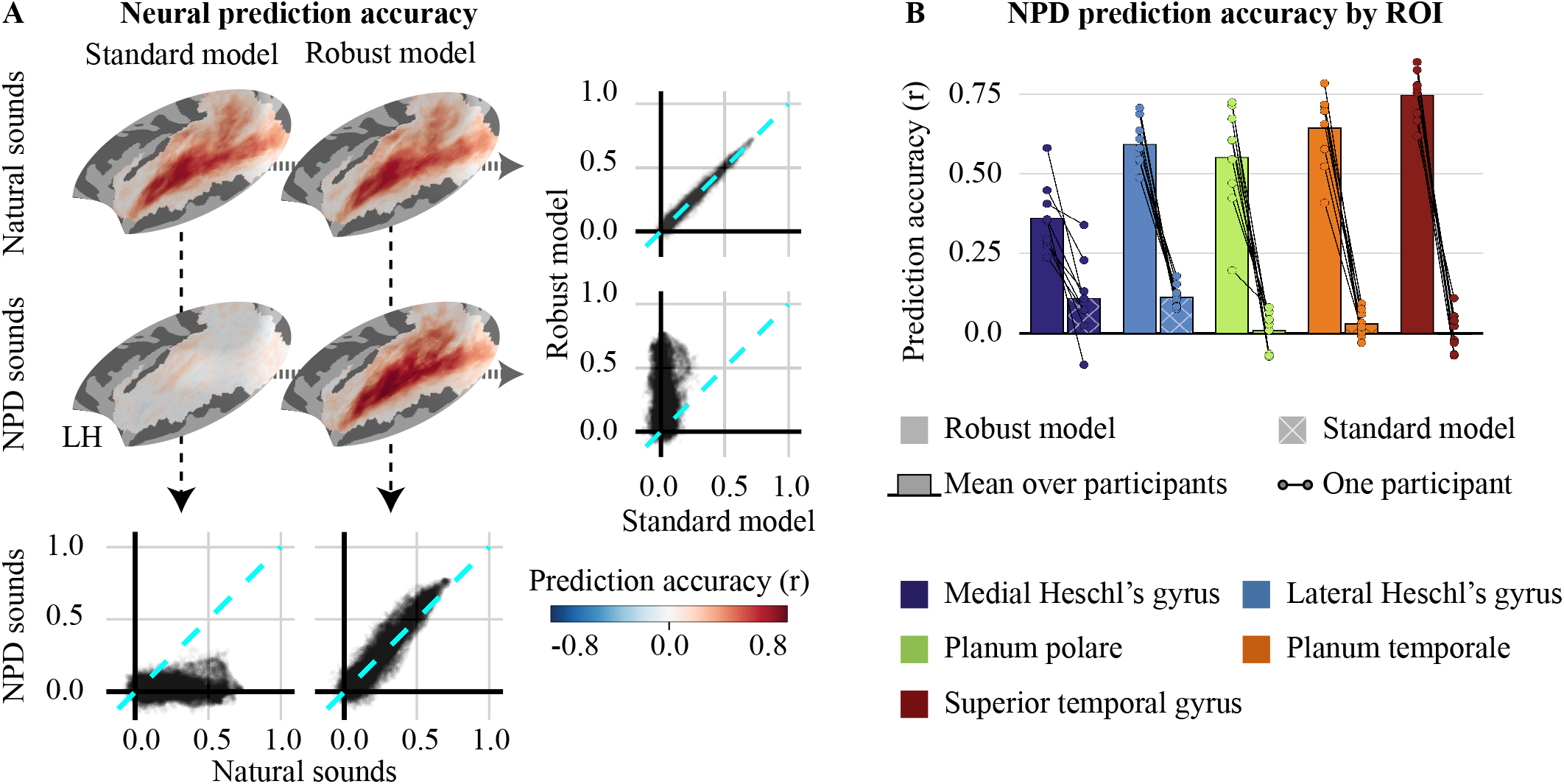
**A** Maps show the neural prediction accuracy for the standard (left column) and robust (right column) models for natural (top row) and NPD (bottom row) sounds. Scatter plots at bottom compare the prediction accuracy between sound sets for the same model. Scatter plots to the right compare the prediction accuracy between models for a given sound set. The prediction accuracy is again virtually indistinguishable for natural sounds. For NPD sounds, the prediction accuracy of the robust model increases slightly, while the standard model accuracy falls to near 0. These maps plot participant-averaged prediction accuracy from the left hemisphere (Supplemental Figure S3A shows right hemisphere maps; Supplemental Figure S3B shows data from individual subjects). **B** Neural prediction accuracy for NPD sounds from robust (solid) and standard (hashed) models across different anatomical regions-of-interest (ROIs). Each circle plots the median prediction accuracy across the voxels in each ROI from both hemispheres in each participant. Lines connect prediction accuracies for standard and robust models from the same participant. Bars plot the average across participants. Prediction accuracy is better for the robust model in every participant and region.

We again found that robust and standard models showed very similar neural prediction accuracy for natural sounds (Fig. 3A), replicating our earlier analyses. For NPD sounds, however, we observed a dramatic difference. The robust model maintained strong neural prediction accuracy for the NPD sounds, even though the models were fit using only natural sounds. In striking contrast, the neural prediction accuracy of the standard model fell to near zero for the NPD sounds. As a consequence, the neural prediction accuracy of NPD sounds was much higher for the robust compared to the standard model in every region of the auditory cortex (Fig. 3B) (Medial Heschl’s gyrus: 0.539 (0.335, 0.822) Lateral Heschl’s gyrus: 0.680 (0.611, 0.744) Planum polare: 0.976 (0.832, 1.000) Planum temporale: 0.916 (0.844, 0.988) Superior temporal gyrus:1.000 (0.904, 1.000)).

## Overlapping cortical responses and divergent model predictions to natural and NPD sounds

The previous results establish the utility of NPD stimuli for model comparison, but it remains possible that NPD sounds evoke voxel response patterns that are highly distinct from those evoked by natural sounds. If so, differences between models may be less relevant for understanding the mechanisms of natural hearing, even when one model shows significantly better prediction performance for NPD sounds. To test this, we correlated the neural response patterns between all pairs of natural and NPD sounds. Correlations between natural and NPD sounds were as high as those within either group (Fig. 4A), demonstrating that NPD sounds produce neural response patterns that overlap with those for natural sounds.

**Fig. 4.**
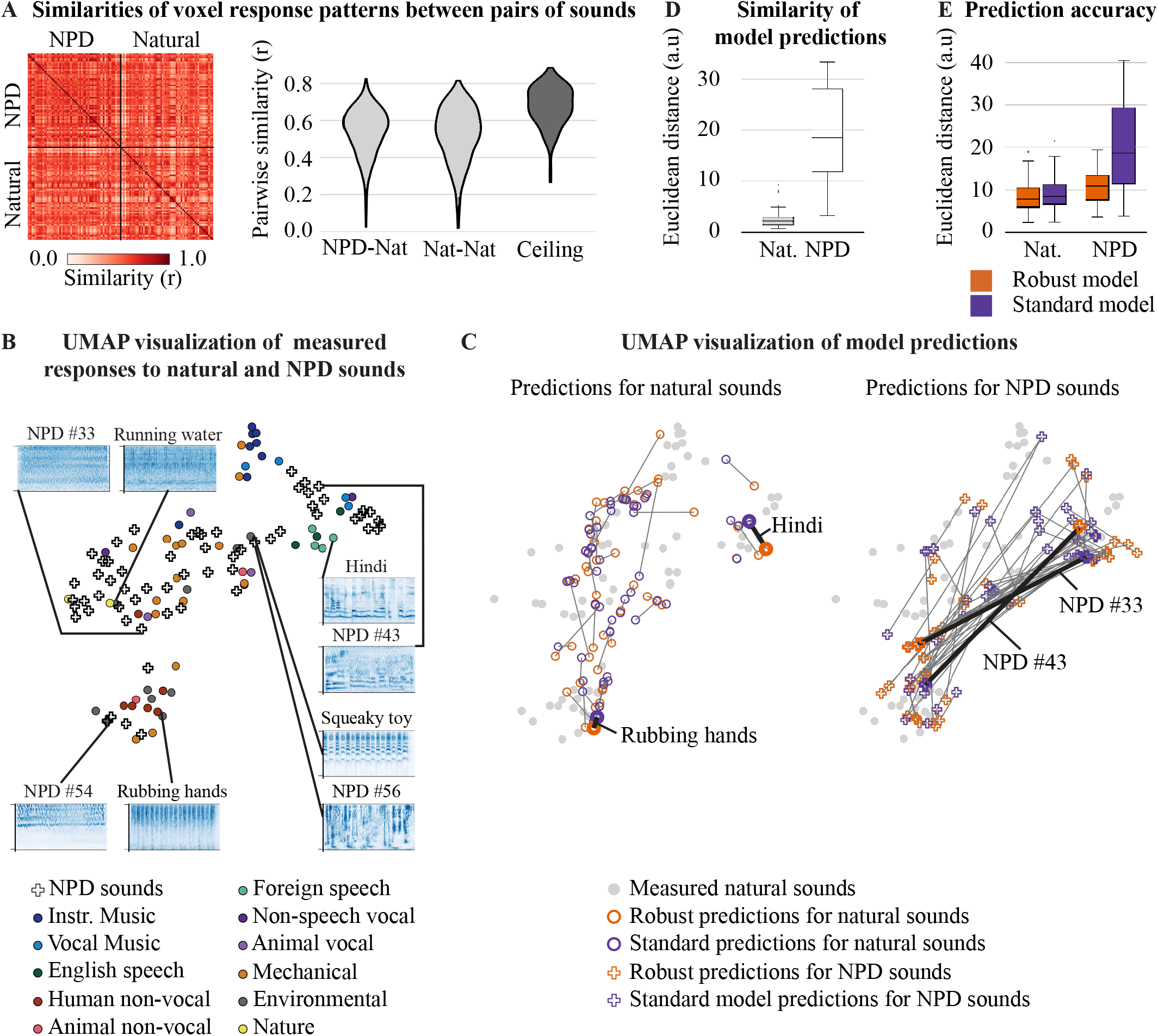
**A** The heatmap shows the similarity (Pearson correlation) between voxel patterns for each pair of NPD and natural sounds. Similarity was computed separately in each participant and then averaged across participants. The violin plots show the distributions of similarity scores when comparing NPD to natural stimuli (leftmost violin) vs. when comparing two different natural stimuli (middle violin), which shows that NPD sounds are about as similar to natural sounds in their response pattern as natural sounds are to themselves. We also plot an estimate of the maximum possible similarity given the reliability of voxel responses to the same sound (rightmost violin). **B** Two-dimensional UMAP embeddings of the voxel responses to natural and NPD sounds. Natural sounds (circles) are colored by sound category; NPD sounds are represented by white crosses. Callouts show cochleagrams for select sounds. **C** UMAP embeddings of standard and robust model predictions for natural (left) and NPD (right) sounds. The model predictions were embedded in the same 2D space used for the measured neural responses in panel **B** so that one can observe the organization of model predictions with the cortical response space. Gray dots in the background show the embeddings for the measured responses to natural sounds (gray; same as panel **B**) for reference. Lines connect predictions of the same stimulus. Note how the robust and standard models make similar predictions for natural sounds in the cortical space but highly divergent predictions for NPD sounds. **D** Similarity of model predictions between robust and standard models for natural (left) vs. NPD sounds (right). Similarity was computed as the Euclidean distance between model predictions in a low-dimensional cortical response space (computed using PCA). Model predictions for natural sounds are similar between the two models, but highly divergent for NPD sounds. **E** Accuracy of the model-predicted response patterns, measured as the Euclidean distance between the measured and model-predicted responses in the low-dimensional cortical space. The robust model is substantially more accurate than the standard model for NPD sounds, but both models are similarly accurate for natural sounds.

To understand how overlapping cortical responses to natural and NPD sounds are organized, we visualized their responses with UMAP after embedding them in a low-dimensional space (spanned by the first 6 principal components, following prior work Norman-Haignere et al. (2015)). We observed interpretable organization based on sound categories and acoustic features that was shared between natural and NPD sounds. For example, synthetic sounds with speech-like structure (e.g., NPD #43) clustered nearby to natural speech sounds. Synthetic sounds with stable spectrotemporal statistics (NPD #33) clustered near natural sound textures with similar statistics (“Running water”) (Fig. 4B). Sounds with high frequency content (NPD #54 and “Rubbing hands”) or sounds with strong temporal modulations (NPD #56 and “Squeaky Toy”) also clustered together irrespective of whether they were natural or synthetic.

We next asked how the predictions from robust and standard models were organized within this neural subspace. We projected the predictions from both models into the shared cortical space, using the same UMAP embedding learned from the measured voxel data. We found that robust and standard models made similar predictions for natural sounds within the shared cortical space, but highly divergent predictions for NPD sounds, evident as highly distinct locations within the shared UMAP space (Fig. 4C). Consistent with this observation, we found that the Euclidean distance between the two models’ predictions within the underlying low-dimensional space was much smaller for NPD compared with natural sounds (Fig. 4D) (distance was measured using the original principal components space, prior to applying UMAP, in order to avoid nonlinear distortions). As expected, the robust model’s predictions for NPD sounds were closer to the measured responses than the standard model’s predictions, while the two models were comparably accurate for natural sounds (Fig. 4E). These results suggest that neural prediction decorrelation was not achieved by pushing responses outside of the distribution for natural sounds, but rather by synthesizing sounds that made divergent predictions within a shared underlying cortical representation space.

## Discussion

Distinct encoding models often make similar neural predictions for complex stimuli, which presents a fundamental challenge for model comparison. We successfully synthesized a sound set that yielded uncorrelated model predictions from adversarially robust and standard DNNs across the entire human auditory cortex in new subjects. NPD sounds evoked responses that overlapped with, and were organized similarly to, those evoked by natural sounds, yet revealed a dramatic difference in neural prediction accuracy that was completely masked when tested using natural sounds alone. Predictions from the standard model were accurate only for the natural sounds, while the robust model remained accurate across the broader NPD stimulus set, suggesting that the robust model can support the design of novel stimuli that evoke specific patterns of neural responses.

### Leveraging stimulus synthesis to test computational sensory models

Sensory neuroscientists have, for decades, characterized neural tuning for simple sensory features using parameterized synthetic stimuli (Hubel and Wiesel 1962; De Valois et al. 1982; Depireux et al. 2001; Woolley et al. 2005). Visual gratings and plaids have been used to reveal tuning for orientation (Hubel and Wiesel 1962) and spatial frequency (De Valois et al. 1982). Tone and noise stimuli have revealed tuning for audio frequency, amplitude modulation, and spectrotemporal modulation (Merzenich and Brugge 1973; Joris et al. 2004; Depireux et al. 2001; Woolley et al. 2005), with modulation tuning properties that appear matched to natural sound statistics (Rodríguez et al. 2010). More sophisticated synthetic stimuli, such as parametric textures (Ziemba et al. 2016) or online closed-loop stimulus optimization (Chambers et al. 2014), have further revealed hierarchical and multidimensional selectivity in sensory cortex. Many cortical regions, however, exhibit complex responses that cannot be easily described based on tuning to parametric features of the input(Theunissen and Elie 2014; Norman-Haignere et al. 2015; Hamilton et al. 2021). As a consequence, many synthetic stimuli produce weak responses in higher-order cortical regions (Norman-Haignere and McDermott 2018; Landemard et al. 2021), such as human non-primary auditory cortex, and models based on tuning for simple stimulus features often struggle to explain responses to more complex stimuli, such as speech (Keshishian et al. 2020; de Heer et al. 2017; DiLiberto et al. 2015). Because of these limitations, it has become increasingly common to directly evaluate models by their ability to predict responses to complex natural stimuli (Yamins et al. 2014; Kell et al. 2018; Tuckute et al. 2023; Naselaris et al. 2011; Khosla et al. 2021).

Our study is part of a growing trend of blending the rigor and control of synthetic stimuli with the predictive power of modern computational models (Bashivan et al. 2019; Ponce et al. 2019; Wang and Ponce 2022; Feather et al. 2023; Walker et al. 2019). For instance, prior studies have designed visual stimuli to maximally drive specific patterns of neural responses (Bashivan et al. 2019; Walker et al. 2019) or to dissociate primary and non-primary auditory cortical responses to natural sounds (Norman-Haignere and McDermott 2018). This previous work highlighted how synthetic stimuli could be utilized with neural encoding models, but did not directly address the challenge of testing which encoding models best predict cortical responses.

We introduce a novel approach that leverages synthetic stimuli to effectively compare models of neural responses. Our work is partially inspired by prior behavioral methods that synthesized stimuli to elicit distinct model decisions (Wang and Simoncelli 2008; Golan et al. 2020), but these methods are not directly applicable to neural population responses, which are high-dimensional and often not directly linked to any single task variable. We address this issue by synthesizing a stimulus set that yields uncorrelated predictions across all neural recordings, simultaneously decoupling the predictions of competing models across the entire population. The synthesized sounds exhibited complex acoustic structures and drove responses comparably to natural sounds in both primary and non-primary auditory cortex (Fig. S4). Yet unlike natural sounds, our synthetic NPD sounds were vastly more effective at model comparison because they were explicitly manipulated to decouple the predictions from competing models.

Why test synthetic stimuli at all when a model accurately captures behavioral or neural responses to natural stimuli? Behavioral studies have shown that DNNs can accurately recognize natural stimuli using features that differ substantially from those of a human observer (Geirhos et al. 2018; Feather et al. 2023), and targeted synthetic stimuli can expose these failures (Goodfellow et al. 2015; Feather et al. 2023). Our results show that the auditory cortex produces similarly structured responses for a wide range of both natural and synthesized sounds. Adversarial training enables robust DNN models to predict this common structure from a shared set of model features, substantially expanding the predictive power of DNNs beyond just natural stimuli. This expansion opens the door to more powerful model-driven experiments, where stimuli are synthesized to achieve targeted scientific or engineering objectives (e.g., model comparison, neural population control).

### The challenge of correlated model predictions

Correlated predictions are pervasive when predicting responses to natural stimuli, which often contain many distinct but correlated features. Speech phonemes, for example, have distinctive spectrotemporal content, and the predictions from phonemes and acoustic models thus often show strong correlations (DiLiberto et al. 2015; Daube et al. 2019). Predictions from DNNs show strong correlations across models with distinct architectures and training techniques (Conwell et al. 2024; Tuckute et al. 2023), as demonstrated here. Correlated predictions are a challenge for model comparison. One approach for addressing correlated predictions is to test whether a model explains unique variance relative to alternative models (de Heer et al. 2017; Brodbeck et al. 2018). However, if two models are maximally correlated across a set of stimuli, as we found to be the case here, then all of the predicted variance will be shared between models and thus not useful for model comparison.

NPD stimuli are designed to reduce correlations in the model predictions for all responses in the sensory population with the goal of facilitating effective model comparison. When measuring neural responses to NPD stimuli, there are three qualitatively distinct possible results. First, one model may show better prediction accuracy than competing models, as observed here for our robust model. Second, both models may show poor prediction accuracy, which provides evidence against both models. Third, both models could show partial prediction accuracy, which provides evidence that both models explain unique parts of the response variance since their predictions are uncorrelated. All three of these outcomes are informative for evaluating model accuracy, unlike cases where models make the same predictions.

### Approaches for decoupling model predictions

Here, we measured and minimized the correlation across sounds separately for each voxel, since model accuracy is often evaluated separately for each neural response in the population (Naselaris et al. 2011; Kell et al. 2018; de Heer et al. 2017; Tuckute et al. 2023). A variant of our method would be to measure the correlation across voxels for each stimulus, as is done in representational similarity analysis (Kriegeskorte et al. 2008), or more generally using other dissimilarity measures between population response patterns (Williams et al. 2021). In practice, we found that minimizing the correlation across stimuli for all voxels yielded sounds that produced highly distinct patterns across voxels. Distinct metrics may thus yield converging results in some cases (Yamins et al. 2014; Conwell et al. 2021; Tuckute et al. 2023), but an interesting future direction would be to explicitly incorporate decorrelated response patterns into the optimization function for NPD stimuli.

Our method also targets high variance in the model predictions. The observed response variance to our NPD stimuli was reliably higher than that for natural stimuli, showing that this approach can be successful (Fig. S4). Variance maximization is particularly useful in cases where there is substantial noise, as with most neural recordings, because it enhances the signal-to-noise ratio.

The NPD method is applicable to any set of differentiable encoding models or neural recording methods. Our method relies on the covariance of model predictions (see 6) and can thus be applied to any number of models. Our loss can be easily modified to incorporate alternative measures of statistical dependency, such as the mutual information between model predictions. In practice, estimating mutual information from small stimulus sets is challenging, and most practical estimators rely on lower-order statistics such as covariance (Kraskov et al. 2004). Our loss function could alternatively be used to select natural stimuli rather than synthesizing new sounds through gradient descent. In our preliminary experiments, we found that synthesis was much more effective than selection at decorrelating model predictions, likely because gradient-based synthesis algorithms are not constrained to a finite set of natural stimuli. Additional constraints could nonetheless be incorporated into our synthesis framework, for example, by biasing the synthesis towards more naturalistic stimuli (e.g., via regularizing constraints or an explicit generative model) (Nguyen et al. 2016; Ponce et al. 2019; Gu et al. 2022; Reilly et al. 2024).

### Benefits of adversarial robustness for neural encoding models

The substantial improvement in neural prediction accuracy observed from adversarial robustness is notable given that standard and robust models shared the same architecture, task (word recognition), and training dataset. These results demonstrate that training DNN models to be robust to adversarial attacks substantially alters the network’s internal representations in a manner that improves neural prediction accuracy across the auditory cortex. These findings resonate with prior behavioral studies suggesting that adversarial training aligns DNN models with human perception (Feather et al. 2023; Gaziv et al. 2023; Tsipras et al. 2019) in both the auditory and visual domains. Parallel physiological work in the visual system supports a link between biological fidelity and adversarial robustness by demonstrating that incorporating biological constraints into standard models improves their prediction accuracy (Dapello et al. 2020) and that perturbations to adversarially trained models also perturb measured neural responses (Guo et al. 2022). Our findings are also consistent with prior vision studies showing similar prediction accuracy of natural images for standard and robust models (Feather et al. 2023; Conwell et al. 2024), even when underlying properties of the representations differ across models (Canatar et al. 2023). Prior findings of similar prediction accuracy for natural stimuli may reflect limitations of the stimulus sets used to compare models, as highlighted by our findings. Our results suggest that adversarial robustness is important for neural population control. Responses to synthetic NPD sounds evoked similar response patterns to natural sounds that overlapped in a shared low-dimensional subspace. Standard and robust DNN models made highly distinct predictions for NPD sounds within this subspace. This observation suggests that each model was attempting to synthesize sounds that produce targeted neural response patterns in the auditory cortex, similar to those that might be observed for natural sounds. The ability of a model to target specific neural response patterns is often referred to as neural population control (Bashivan et al. 2019; Gu et al. 2022; Walker et al. 2019; Wang and Ponce 2022), and to our knowledge, has not been demonstrated in the auditory cortex. In our setting, neural population control is more challenging because the models must synthesize stimuli for which a competing model disagrees about the predicted neural pattern, creating a form of competitive population control. Because of this competition, the measured neural data has an opportunity to adjudicate which model is more effective at population control.

Standard models may have difficulty controlling neural population activity because they only make accurate predictions for the small portion of stimulus space occupied by natural stimuli. While prior studies have reported effective population control in the visual system using standard (non-robust) DNN models, these studies utilized regularization techniques to constrain the synthesized stimuli (e.g., smoothness constraints or generative models) (Ponce et al. 2019; Walker et al. 2019). A recent study using a standard DNN to synthesize sounds targeting speech- and music-selective regions of human auditory cortex similarly required smoothness constraints during synthesis and achieved only partial activation of the targeted regions relative to natural sounds (Xing et al. 2026). If a DNN model accurately describes the underlying neural computations, no additional regularization constraints should be necessary to control neural population activity. Our findings suggest that adversarial training expands the space of stimuli that can be accurately predicted by DNN models to an extent that enables neural population control in human auditory cortex without any additional regularization constraints. This naturally paves the way for future experiments using adversarially trained audio models to perform other types of neural population control experiments and investigate auditory representations.

## Methods

### Overview of Experiments

#### NH2015 and B2021 experiments: fMRI responses to natural sounds

We used fMRI data from two prior experiments that measured responses to a diverse set of natural sounds, which we label NH2015 (Norman-Haignere et al. 2015) and B2021 (Boebinger et al. 2021). The data from these experiments were used to measure neural prediction accuracy and assess the similarity of the model predictions for natural sounds between robust and standard DNNs (Fig. 1). Encoding models fit to these data were also used to synthesize NPD sounds. The data from these experiments have been described in detail in prior studies (Norman-Haignere et al. 2015; Boebinger et al. 2021), but we repeat key methods here so that the reader does not need to consult prior papers. These studies were approved by the Massachusetts Institute of Technology Committee on the Use of Humans as Experimental Subjects, as reported in the original publications.

#### S2026 experiment: fMRI responses to NPD and natural sounds

We conducted a new experiment measuring fMRI responses to synthesized NPD sounds as well as a reference set of natural sounds for comparison. These data were used to measure and compare the neural prediction accuracy of the standard and robust DNN models using both NPD and natural sounds (Fig. 3), as well as to understand how voxel response patterns and model predictions for NPD and natural sounds are organized in the auditory cortex (Fig. 4). The study was approved by the University of Rochester Research Subjects Review Board.

### Participants

NH2015 tested 10 participants (4 male, 6 female, ages 19-27). B2021 tested 20 participants (6 male, 14 female, ages 18-34), half of whom were trained musicians and half of whom had little musical training. A prior study (Boebinger et al. 2021) found that fMRI responses in the auditory cortex were very similar between the two groups, and data were therefore pooled across both groups. S2026 tested 9 participants (5 male, 4 female, ages 18-35). There was no overlap in the participants tested in each experiment. All participants had self-reported normal hearing. Audiometry was conducted for participants in B2021 and S2026, and all participants had normal hearing thresholds (below 25 dB HL for octave frequencies 250 Hz to 8 kHz). All participants were native English speakers.

### Data collection and preprocessing

Data for the NH2015 and B2021 were collected on a 3T Siemens Trio scanner using a 32-channel head coil at the Athinoula A. Martinos Imaging Center of the McGovern Institute for Brain Research at MIT. Data for S2026 were collected on a 3T Siemens Prisma Scanner with a 64-channel head coil at the Center for Advanced Brain Imaging and Neurophysiology (CABIN) at the University of Rochester. Details about the parameters of the MRI scan for NH2015 and B2021 are reported in (Norman-Haignere et al. 2015; Boebinger et al. 2021), respectively. The parameters for S2026 are included in supplemental figure S6.

Experiment NH2015 and B2021 measured responses to a set of 165 natural sounds, each 2-seconds in duration. The sounds were selected to include many of the most commonly heard and recognizable sounds encountered in daily life (Norman-Haignere et al. 2015). The sounds were grouped into 11 different sound categories, determined based on participant ratings from the NH2015 study: instrumental music, music containing vocals, English speech, foreign speech, non-speech human vocalizations (e.g., laughter), animal vocalizations, human non-vocal sounds, animal non-vocal sounds, nature sounds, mechanical sounds, and environmental sounds. Foreign speech included speech from 7 different languages (German, French, Italian, Russian, Hindi, Spanish, Chinese), most of which were unfamiliar to the participants, who were all native English speakers. B2021 included an additional set of 27 non-Western music and drumming clips, but for simplicity, we focused on just the 165 sounds that were common between the two studies. S2026 measured responses to 120 natural sounds, which were a subset of the 165 sounds used in NH2015 and B2021, plus 60 NPD sounds (see description below). Sounds were RMS-normalized and presented via Sensimetrics S14 earphones.

Sparse scanning was used to prevent scanner noise from interfering with participants’ ability to hear the sounds. Each fMRI scan acquisition was 1 second in duration, after which a single 2-second sound was presented during a 2.4-second gap between scan acquisitions (with a 200 ms silent buffer before and after each sound) (Fig. S6). Each sound was repeated multiple times in a short block (5 sounds per block in NH2015, 3 sounds per block in B2021/S2026) because we have found that this enhances the reliability of the measurements. To ensure participants were attending to the sounds, they performed a simple task in which they detected a change in loudness. Specifically, one of the sounds in each block was quieter than the other sounds (by 7 dB in NH2015 and by 12 dB in B2021/S2026), and participants were instructed to press a button for the quieter sound.

We collected data from participants over multiple scanning sessions to make it possible to collect additional data in each subject. In NH2015, participants completed 2-3 scanning sessions (≈2 hours per session; 1 block per sound per session). In B2021, participants completed 3 scanning sessions (≈2 hours per session, 2 blocks/session). S2026 used a mixed design where seven participants completed 2 scanning sessions (≈75 minutes per session, 1 block/session) and two participants completed 6 sessions. This mixed design, which we have successfully used in prior work (Norman-Haignere and McDermott 2018), makes it possible to compute group-level statistics by testing many participants, each with a smaller amount of data, as well as compute reliable brain maps from a smaller number of individual participants with many scans per subject.

Functional volumes were motion-corrected and aligned to the anatomical volumes. Volume data were then resampled to the cortical surface and lightly smoothed using a 3 mm full-width-at-half-maximum (FWHM) kernel. We have previously found that smoothing data using a 3 mm kernel improves prediction of unsmoothed data, suggesting that smoothing at this scale mostly suppresses unreliable noise (Norman-Haignere et al. 2015). Preprocessing was implemented using a mixture of FSL (NH2015, B2021), fMRIPrep (S2026), and Freesurfer (all experiments). S2026 used fMRIPrep’s default parameters (version 25.1.4). Code detailing the preprocessing steps will be released upon publication, along with original (defaced) and preprocessed data. For B2021 & S2026, a general linear model was used to estimate the response of each voxel to each stimulus block (convolving each block with a standard hemodynamic response function). For NH2015, a simpler signal averaging procedure was used because it was found to be more reliable for the longer blocks tested in this experiment (the 2nd through 5th acquisition after each stimulus block was averaged).

We selected voxels with a reliable response to sound. For group-level maps (Fig. 1B&D, Fig. 2C, Fig. 3A), we selected voxels at the group level by measuring reliability across splits of subjects. Specifically, we selected voxels using the following criterion:

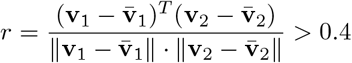

where **v**_1_ and **v**_2_ are the average responses for a single voxel in standardized coordinates from two non-overlapping sets of 15 participants across both NH2015 and B2021. We used the same voxels in Fig. 3A even though that figure shows S2026 data for comparability with earlier results.

Analyses that quantified effects or response patterns at the individual subject level (Fig. 1C&E, Fig. 3B, Fig. 4) were based on subject-specific reliability computed across repetitions. Specifically, analyses used voxels within the same broad anatomical region with test-retest reliability greater than 0.4, computed using the above equation. For S2026 participants with 6 repetitions, even and odd repetitions were averaged before computing the test-retest reliability.

### DNN encoding models

Voxel-wise encoding models were derived from deep neural networks (DNNs) previously shown to yield good fMRI prediction accuracy in the human auditory cortex (Tuckute et al. 2023). The full details of the model training are available in (Feather et al. 2023), but we repeat key details here to minimize the need to consult external sources. DNNs were trained to classify spoken words at the center of a 2-second clip using a standard cross-entropy loss (794-way classification consisting of 793 words plus a single “null” class representing the speech absent condition). Background sounds were added to the speech for most samples of model training to make the recognition task more challenging (SNR ranged from -10 to 10 dB). The input to the DNN was a cochleagram representation, computed by convolving each sound with a collection of bandpass filters (50-10000 Hz on an ERB scale (Glasberg and Moore 1990)), computing the envelope from the output of each bandpass filter, applying compression (compression factor = 0.3), and then downsampling (from 20000 to 200 Hz). The result is a 2D time-frequency “image” of sound that coarsely mimics the information present in the auditory nerve (McDermott and Simoncelli 2011). Cochleagrams were RMS-normalized and fed as input into a deep neural network with a “ResNet50” architecture (He et al. 2016). The full details of the architecture are provided in (Feather et al. 2023) and in Supplemental Table ST1.

Our standard and robust DNNs were identical, except that the robust DNN underwent additional training to make it robust to adversarial attacks (Madry et al. 2018). The approach is to generate a small perturbation to a stimulus that causes the model to misclassify that stimulus. The perturbed stimulus is then fed back into the model, and the model is trained to correctly classify that stimulus. A perturbation was generated by performing gradient descent on the cochleagram representation to maximize (rather than minimize) the cross-entropy loss. The perturbation was constrained to be small by setting the *l*^2^ norm of the perturbation to 1. We then performed gradient descent on the model parameters to minimize the cross-entropy loss. Additional details of the training procedure can be found in (Feather et al. 2023).

We fit voxel-wise encoding models by regressing the activations of all units from a single DNN layer onto the response of each voxel. Ridge regression was used to regularize the model and prevent overfitting. To reduce computational demands, the regression analysis was performed using principal components of the activation matrix (preserving the variance of each component, which is important for regularization). The number of principal components was set equal to the number of training sounds, and thus no information was lost when converting to principal components. For each voxel, we used activations from the layer that best predicted that voxel’s response (either layer1, layer2, layer3, layer4, or avgpool). Consequently, we selected two hyperparameters for each voxel and model: the DNN layer and the ridge penalty parameter (from a broad array of 500 values evenly spaced on a logarithmic scale between 1 × 10^−8^ and 1 × 10^8^). We used nested cross-validation (5 outer folds, 5 inner folds) to select both hyperparameters: the inner loop was used to choose the ridge penalty for each layer, and the outer loop was used to generate cross-validated predictions for selecting the best layer. Performance of the final model was evaluated on a separate, held-out set of natural sounds.

We measured prediction accuracy by computing the Pearson correlation between the measured and model-predicted voxel responses across 55 test sounds not used to fit the encoding models (Fig. 1A). Prediction accuracy was always measured in voxels from the individual subjects (not group-averaged data). For group-level maps, we then averaged the individual subject prediction accuracies across participants in standardized coordinates. We also measured the similarity of the model predictions by correlating the predictions across robust and standard models (Fig. 1B) (using the best-predicting layer for each model). Since the predictions were computed by regression against noisy neural responses, voxels that were less reliable also had less correlated predictions. To determine the maximum possible correlation we could expect between two models, given the reliability of the voxel responses, we fit the same DNN model (e.g., just the standard DNN) using two non-overlapping sets of sounds and measured the correlation of the predictions in a third, left-out sound set. Since in this case, the DNN network is identical, any difference in the model predictions can only be due to variability in the voxel responses. We then compared this “Same DNN” correlation (standard vs. standard, robust vs. robust) with an “Across DNN” correlation, computed by correlating the predictions between standard and robust DNNs.

### Neural prediction decorrelation

We give a complete description of our method for synthesizing stimuli that decorrelate neural predictions. We repeat some of the content from the Results so that this section is comprehensive. Nothing about the method is specific to sounds, fMRI, or DNN models, and the method is applicable to any number of models. We assume we have a set of *m* ∈ 1 … *M* models that can make predictions 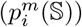 for any set of stimuli (S) for all channels (*i* ∈ 1 … *N* ) in a neural population. The model predictions can be any differentiable function that maps from stimuli to the predicted neural responses. For a given stimulus set, S, we compute an *M* × *M* covariance matrix **Φ**_*i*_(S) (Equation 1) across stimuli between the predictions of all models. The off-diagonal elements of this matrix contain the covariance (Equation 3) between the predictions for all pairs of models. The on-diagonal elements of the matrix contain the variances (Equation 2) of the predictions. NPD stimuli are synthesized so that the empirically measured covariance matrix **Φ**_*i*_(S) matches a target covariance matrix (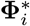, Equation 4).

The off-diagonal elements of the target covariance matrix are set to 0 to encourage uncorrelated predictions, and the on-diagonal elements are set so as to encourage high variance. Here we set the variance targets to be a multiple (*α* = 5) of the variance for a reference set of natural sounds, which we empirically found led to uncorrelated, high-variance predictions (S5, as desired).

We synthesized NPD sounds by minimizing the loss shown in Equation 5 with respect to S. The sounds were 2-seconds and initialized with unstructured Gaussian noise (*i*.*i*.*d*. samples from N (0, .01) at a sampling rate of 20000 Hz), and we performed gradient descent on the sounds using the Adam optimizer (Kingma and Ba 2017) for 10000 iterations to minimize the loss. The learning rate followed a cosine annealing schedule with warm restarts every 1000 iterations, with an initial learning rate of 1 × 10^−3^ and reaching a minimum learning rate of 1 × 10^−5^. During optimization, the sounds were windowed using linear ramps, highpass filtered at 50 Hz, and RMS normalized to 0.1 (these three operations were applied every 100 iterations). Because the effects of repeated windowing will compound over optimization, the duration of the ramps varied over training to achieve symmetric, approximately 100 ms linear ramps by the final iteration. Optimization was done in PyTorch (Ansel et al. 2024). We synthesized 60 NPD sounds using this procedure.

We synthesized sounds using encoding models fit to voxels from a single experiment (NH2015) and then evaluated whether the synthesized sounds produced decorrelated predictions using encoding models fit to voxels from a separate experiment (B2021) with entirely new and non-overlapping participants (Fig. 2). We then measured responses to these synthesized sounds in a new experiment (S2026) along with a set of 120 natural sounds, which we used to measure neural prediction accuracy (Fig. 3).

### Measuring similarity of voxel response patterns

To characterize the similarity between voxel response patterns to natural and NPD sounds, we correlated the voxel response patterns for every pair of natural and NPD sounds (Fig. 4A). The correlation was computed separately for each individual participant and then averaged across participants. We computed a noise ceiling by measuring the Spearman-Brown (Spearman 1904) corrected correlation between repeated presentations of the same sound.

To visualize the organization of response patterns, we applied Uniform Manifold Approximation and Projection (UMAP; McInnes et al. 2020) (*k* = 5 nearest neighbors, minimum distance = 0.25, Euclidean metric) (Fig. 4B). Because voxel responses are noisy, voxel responses were reduced in dimensionality by projecting onto the top 6 principal components, following prior work (Norman-Haignere et al. 2015). Results were robust to the choice of the number of principal components. A single UMAP embedding was fit using principal-component-projected fMRI responses for natural and NPD sounds. Model predictions for natural and NPD sounds were subsequently mapped into this fixed embedding space (Fig. 4C, left and right, respectively), so that we could observe how model predictions were organized in the neural space. We measured the similarity of standard vs. robust model predictions as the Euclidean distance in our lower-dimensional component space (top 6 principal components). We measured the accuracy of model predictions as the distance in component space between the measured and model-predicted responses, separately for the robust and standard models. Both analyses were performed separately for natural and NPD sounds. Fig. 4D&E report the median across stimuli; 95% confidence intervals were estimated by nonparametric bootstrap (5,000 resamples with replacement).

### Standard statistical quantifications and region-of-interest analyses

We quantified prediction accuracy using a set of standard regions of interest (ROI) (medial and lateral Heschl’s gyrus, planum polare, planum temporale, and the superior temporal gyrus) (Norman-Haignere et al. 2025; Destrieux et al. 2010) (Fig. 1). For each ROI, we measured the median Pearson correlation across voxels from each participant, and then averaged these quantities across participants. We used a simple selectivity metric to quantify the similarity (or dissimilarity) of the neural prediction accuracy between standard and robust models in each ROI (Fig. 1D):

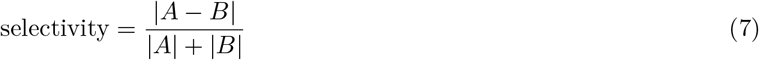

where *A* and *B* are the average Pearson correlation for standard and robust models, respectively. Values near 0 indicate similar accuracy, while values near 1 indicate maximally different accuracy. We computed confidence intervals (central 95%) by bootstrapping this selectivity metric across participants (1000 samples).

## Code and data availability

Code implementing the synthesis procedure and analyses is available at https://github.com/dskrill/neural-prediction-decorrelation. Data and trained models are available at https://doi.org/10.5281/zenodo.22038940.

## Acknowledgements

This study was supported by the National Institutes of Health (NIDCD R00-DC018051, NIDCD R01-DC020960) and the National Science Foundation (2342908). This work was partially supported by the Center for Computational Neuroscience at the Flatiron Institute of the Simons Foundation. Experiments were performed on the Flatiron Institute’s high-performance computing cluster.

## Author contributions

D.S and S.V.N.-H. collected data for the experiments. J.F. contributed the trained deep neural network models. D.S. performed the encoding model fitting, sound synthesis, and analyses with mentorship and guidance from S.V.N.-H. and J.F. D.S, J.F., and S.V.N.-H. collectively wrote the manuscript.

## Supplement

**Fig. S1.**
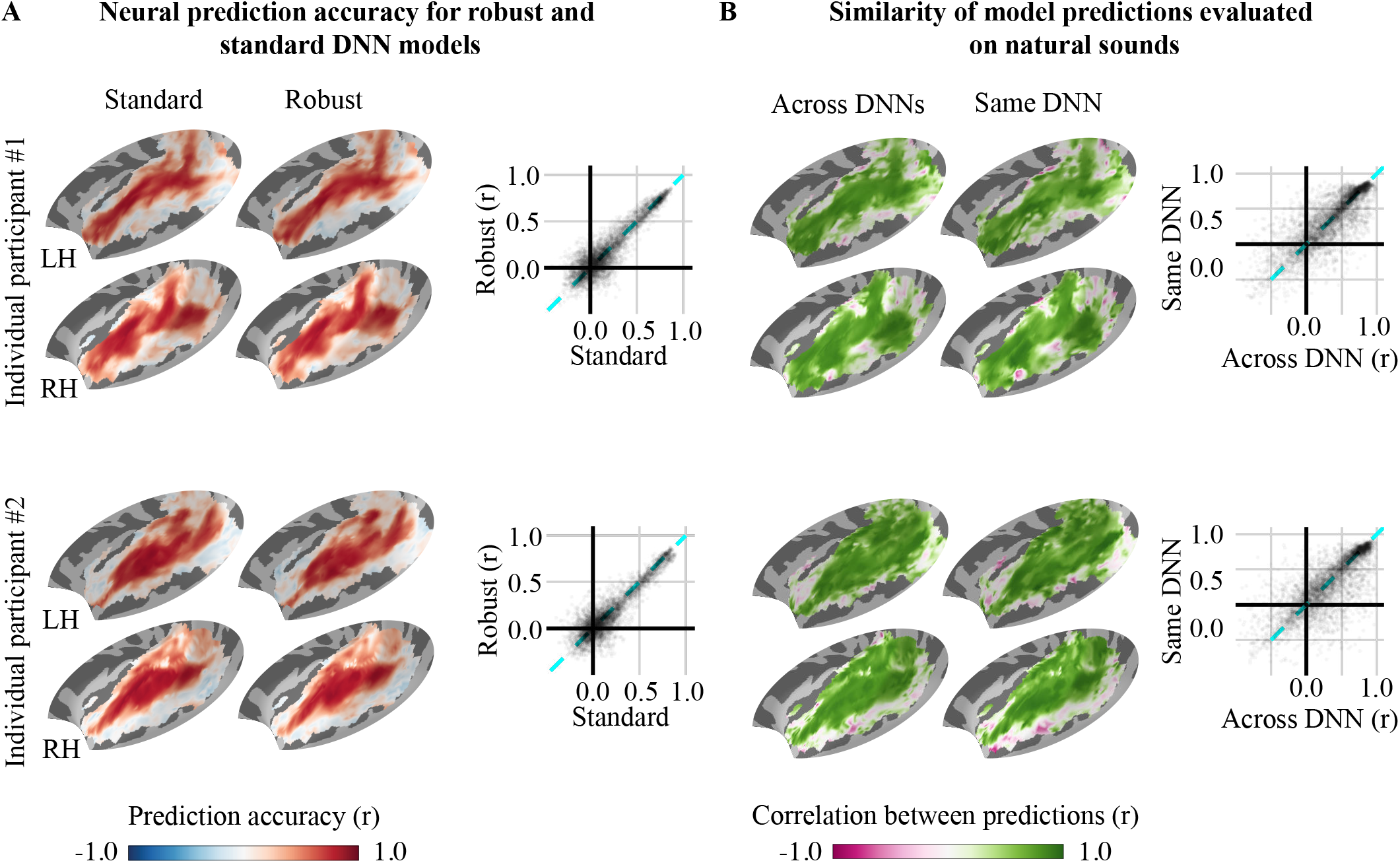
**A** Prediction accuracy (Pearson correlation) for standard (left) and adversarially robust (right) models for individual participants. The scatter plot compares the prediction accuracies of the standard (x-axis) and adversarially robust (y-axis) models for all voxels analyzed. **B** Similarity of the adversarially robust and standard model predictions obtained by correlating the model predictions across models for each voxel (“Across DNNs”, left). The models shown here use different DNNs and were fit to neural responses using non-overlapping training stimuli. This allows us to plot a measure of the maximum possible correlation computed by correlating the predictions from two models with the same DNN (e.g., standard vs. standard, robust vs. robust) fit to neural responses using non-overlapping stimulus sets (“Same DNN”, right). The scatter plot compares the Across DNN correlations (x-axis) with the Same DNN correlations (y-axis) for all analyzed voxels.

**Fig. S2.**
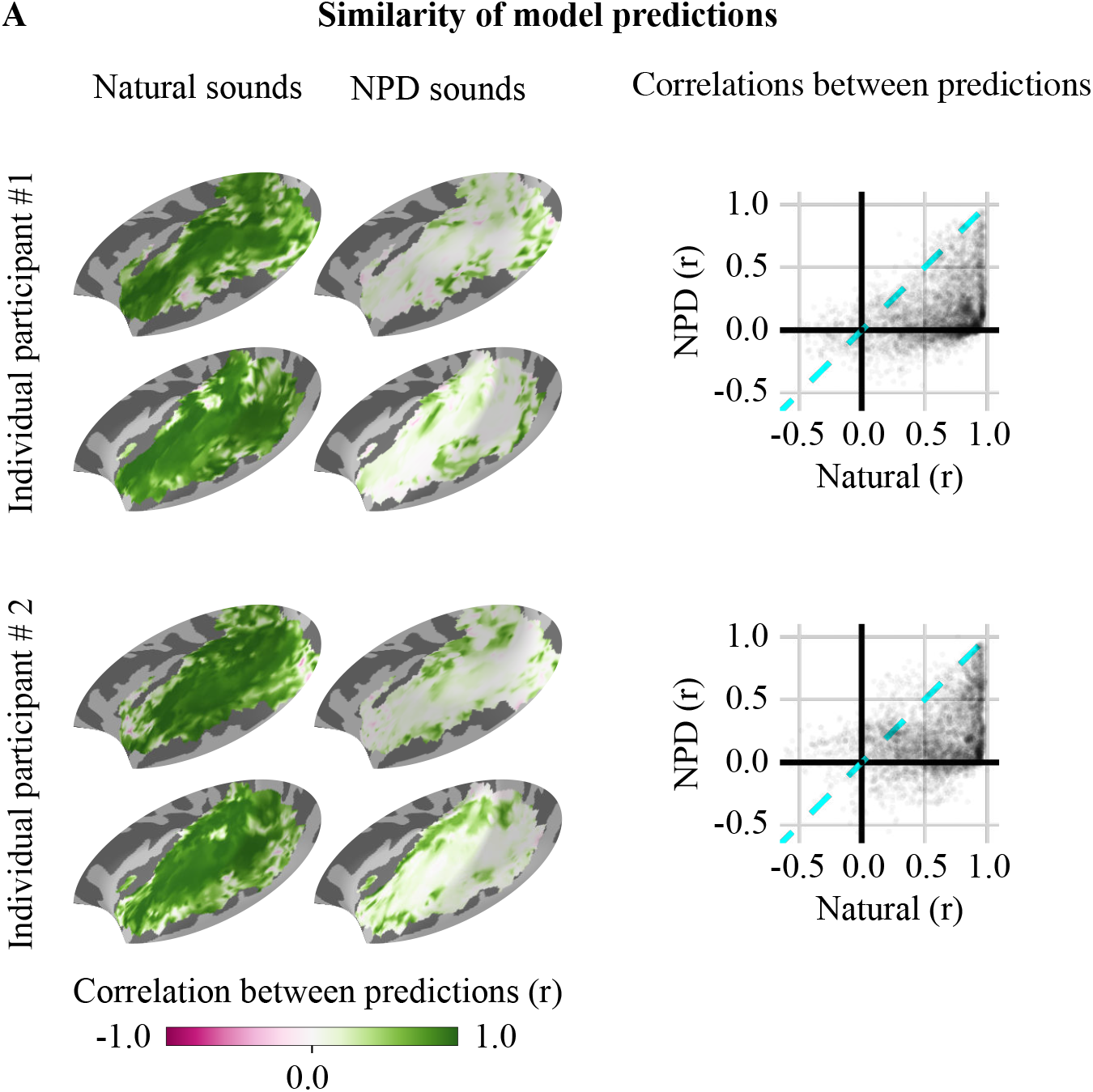
**A** Similarity of model predictions for individual participants. Correlations between model predictions are shown for natural (left map) and NPD (right map) stimuli. The scatter plot compares correlations between model predictions for the natural sounds (x-axis) and NPD (y-axis) sounds.

**Fig. S3.**
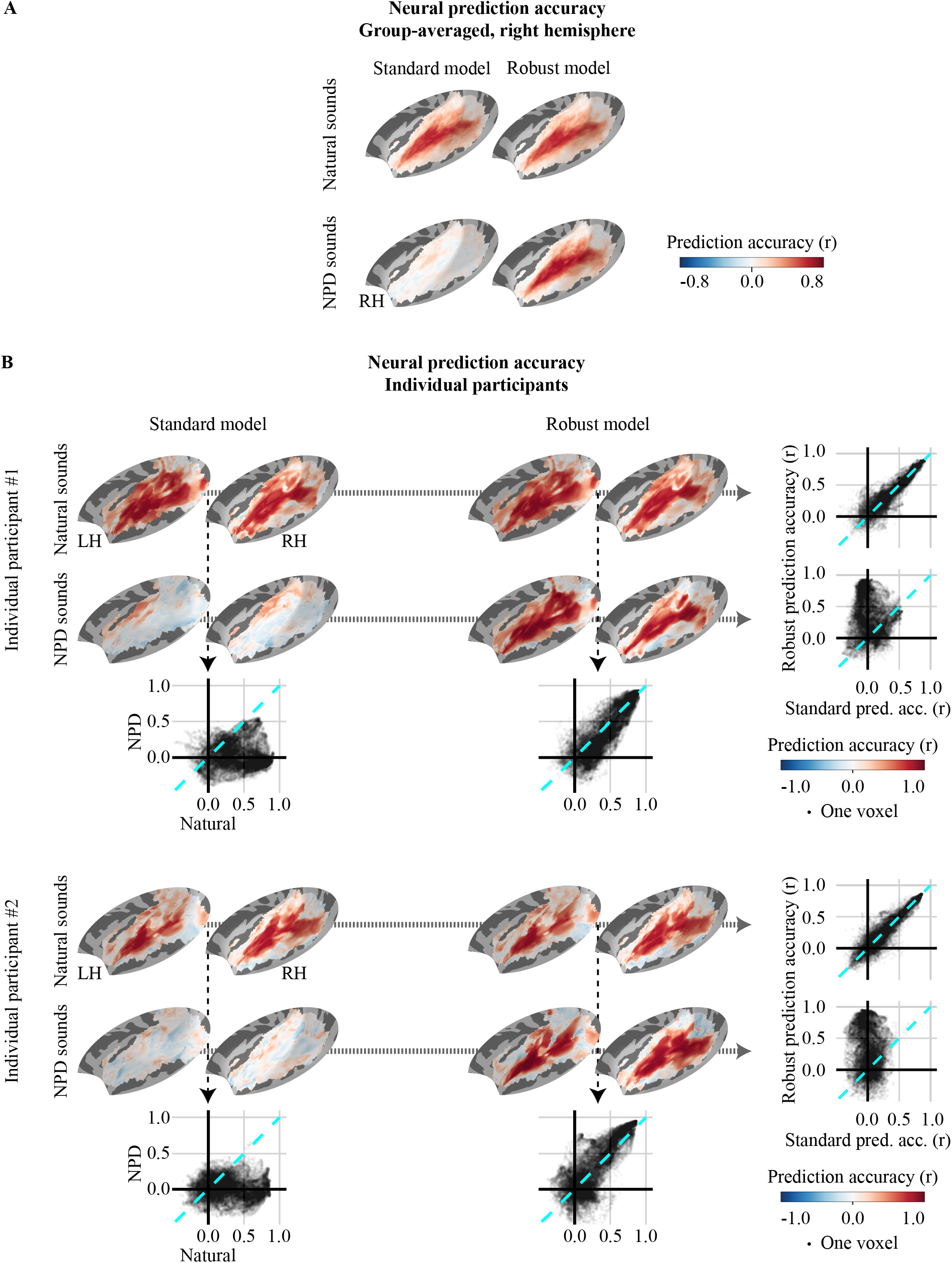
**A** Group-averaged neural prediction accuracy for natural and NPD sounds in the right hemisphere. Prediction accuracy is shown for the standard (left column) and robust (right column) models for natural (top row) and NPD (bottom row) sounds. **B** Prediction accuracy for natural and NPD sounds for two individual participants. Prediction accuracy of the standard (left column) and adversarially robust (right column) models for natural (top rows) and NPD (bottom rows) sounds. The bottom scatter plots compare the prediction accuracy for natural and NPD sounds within a model (e.g., standard vs. standard). The scatter plots on the right compare across models (e.g., adversarially robust vs. standard predictions of natural (top) and NPD (bottom) sounds).

**Fig. S4.**
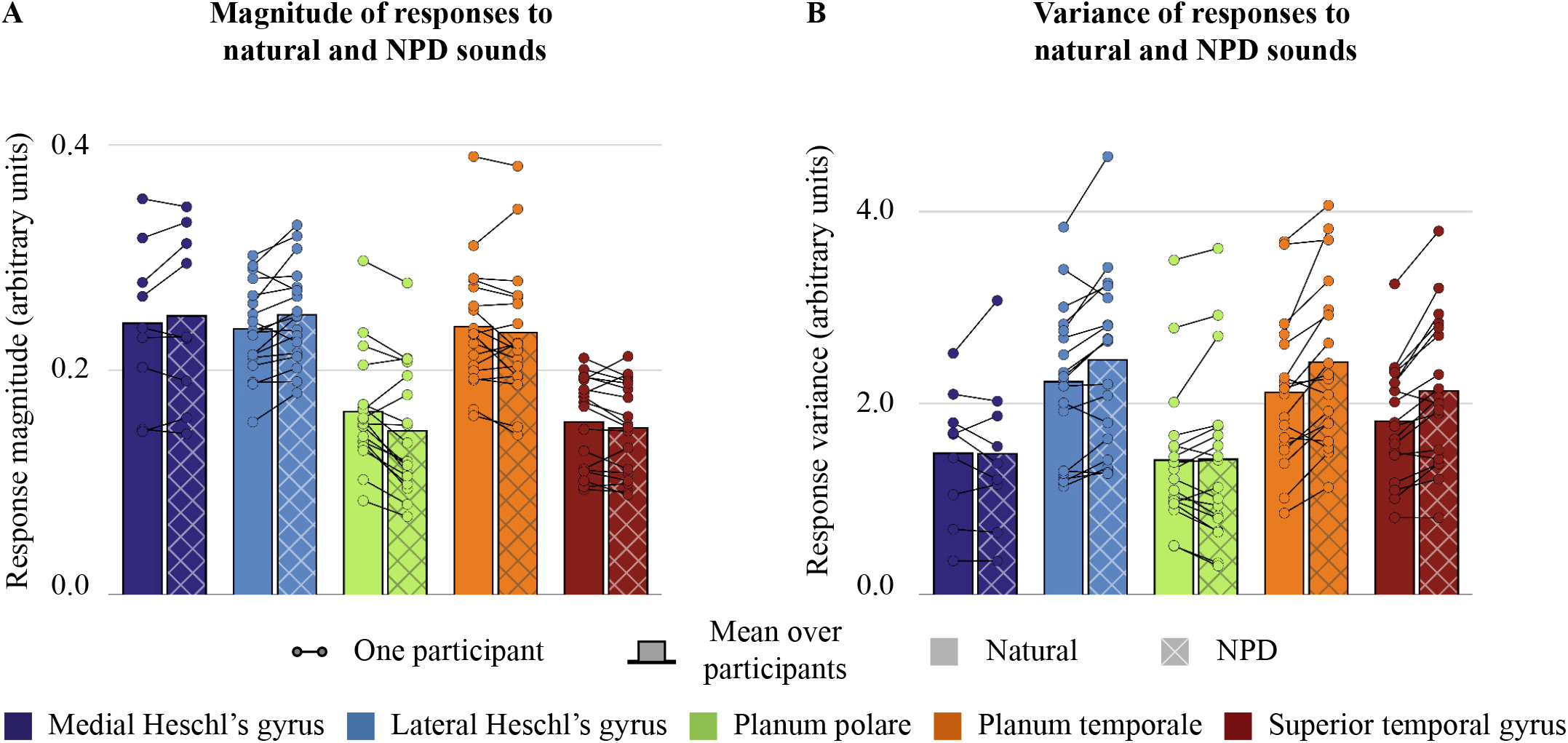
**A** Magnitude of measured responses to natural and NPD sounds in each ROI. Circles show the median response across voxels for each ROI for a single participant. Bar plots show the mean across participants for natural sounds (solid) and NPD sounds (cross-hatched). **B** Variance of measured responses to natural and NPD sounds in each ROI. Same format as panel A.

**Table ST1.**
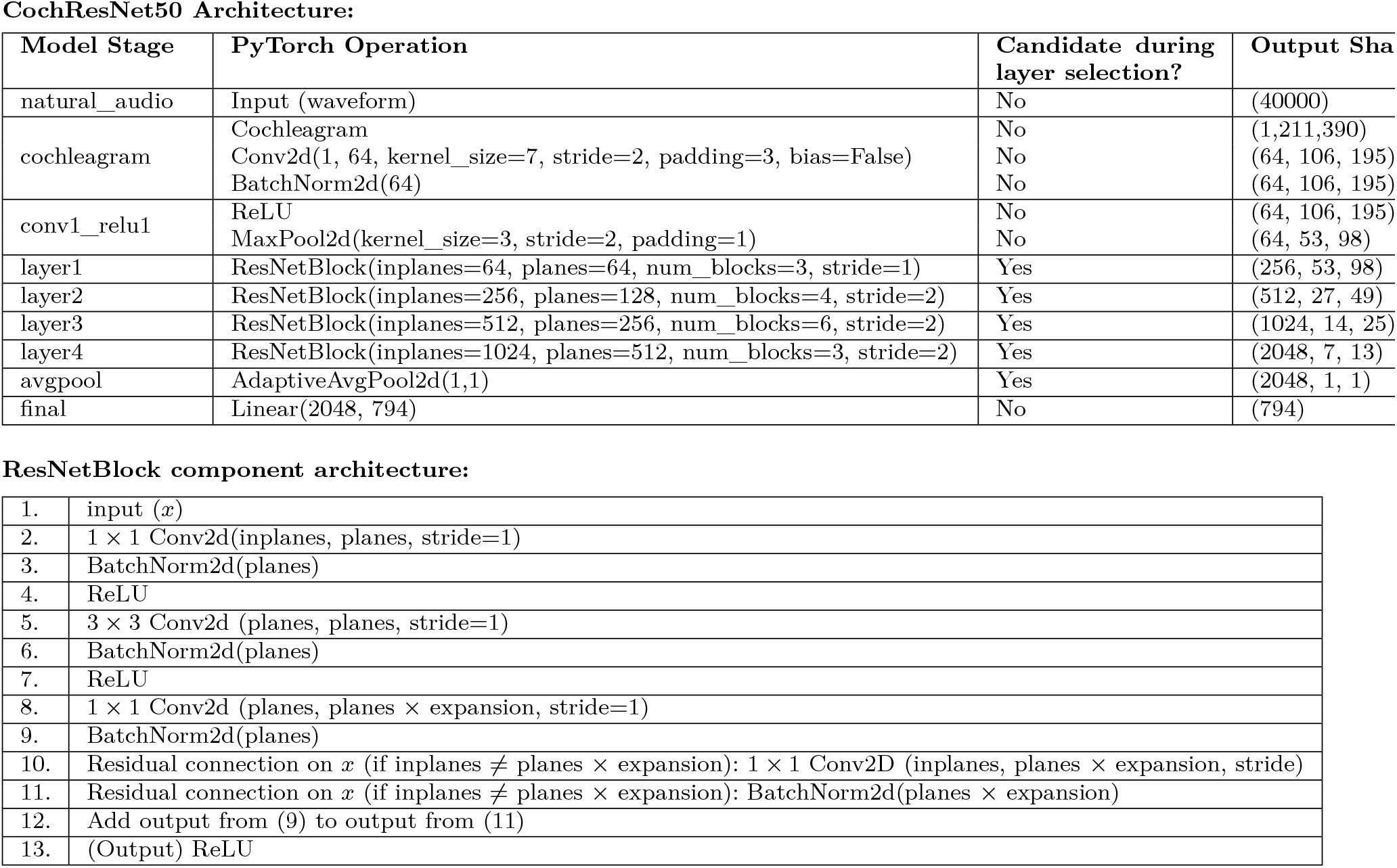
CochResnet50 architecture. The DNN takes a 2-dimensional (time × frequency) cochleagram representation of sound as input and processes it using a ResNet50 backbone architecture (He et al. 2016). Each ResNetBlock consists of num_blocks copies of the ResNetBlock component architecture above, stacked together to form a single ResNetBlock. The expansion factor was set to four for all layers. Only the outputs of layer1, layer2, layer3, and layer4 were considered during layer selection as part of the encoding model fitting (as indicated in the “Candidate during layer selection?” column; see Methods for details).

**Fig. S5.**
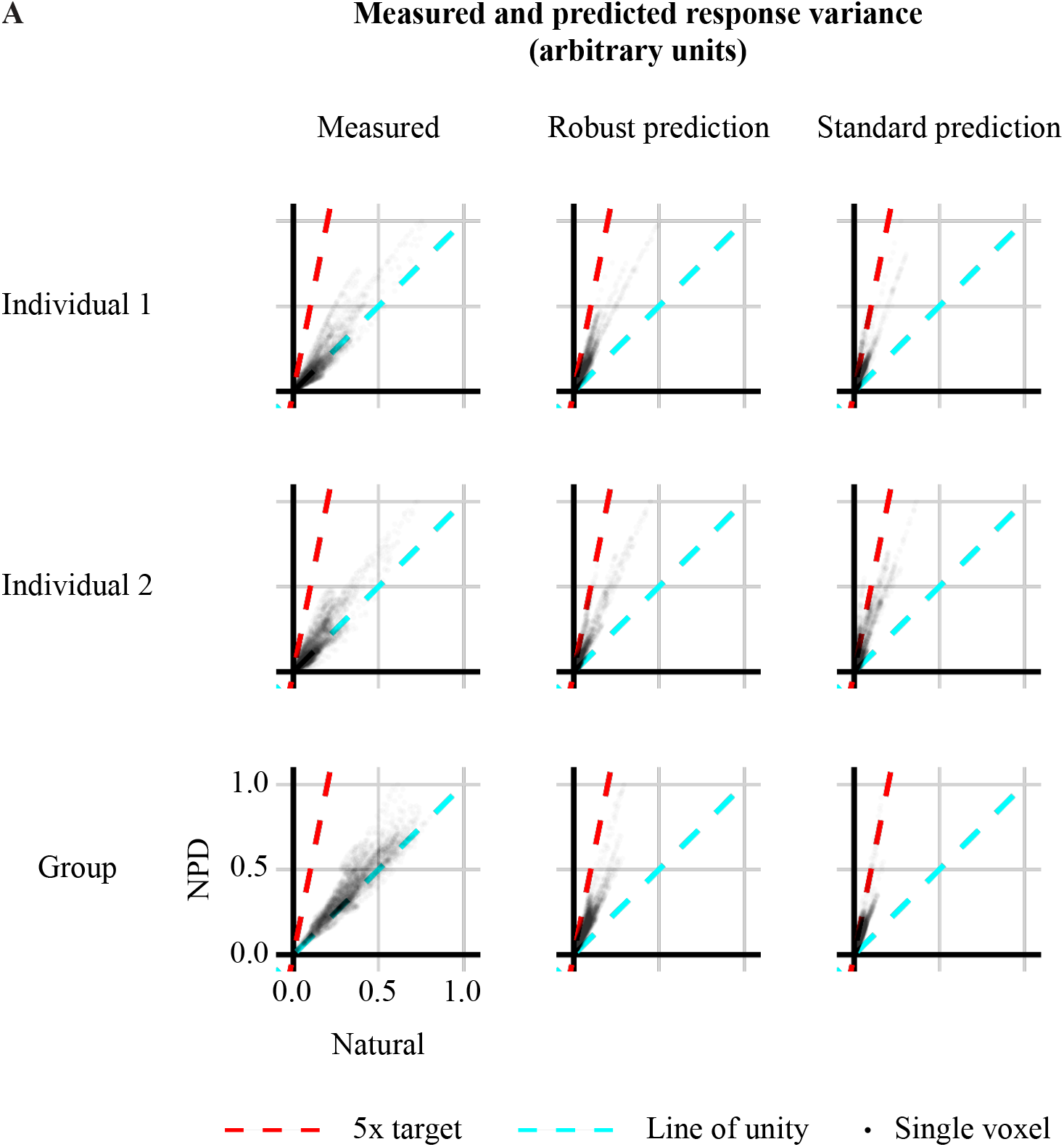
**A** Voxel-wise variances of measured (left column) and predicted (middle and right columns) responses, for two individual participants (top and middle rows) and at the group level (bottom row). The 5× variance target is shown in each plot. We observe that NPD sounds tend to produce higher variance responses than natural sounds, as desired. This difference is smaller than the difference in predicted variance, plausibly because measured neural responses include noise that is independent of the stimuli.

**Fig. S6.**
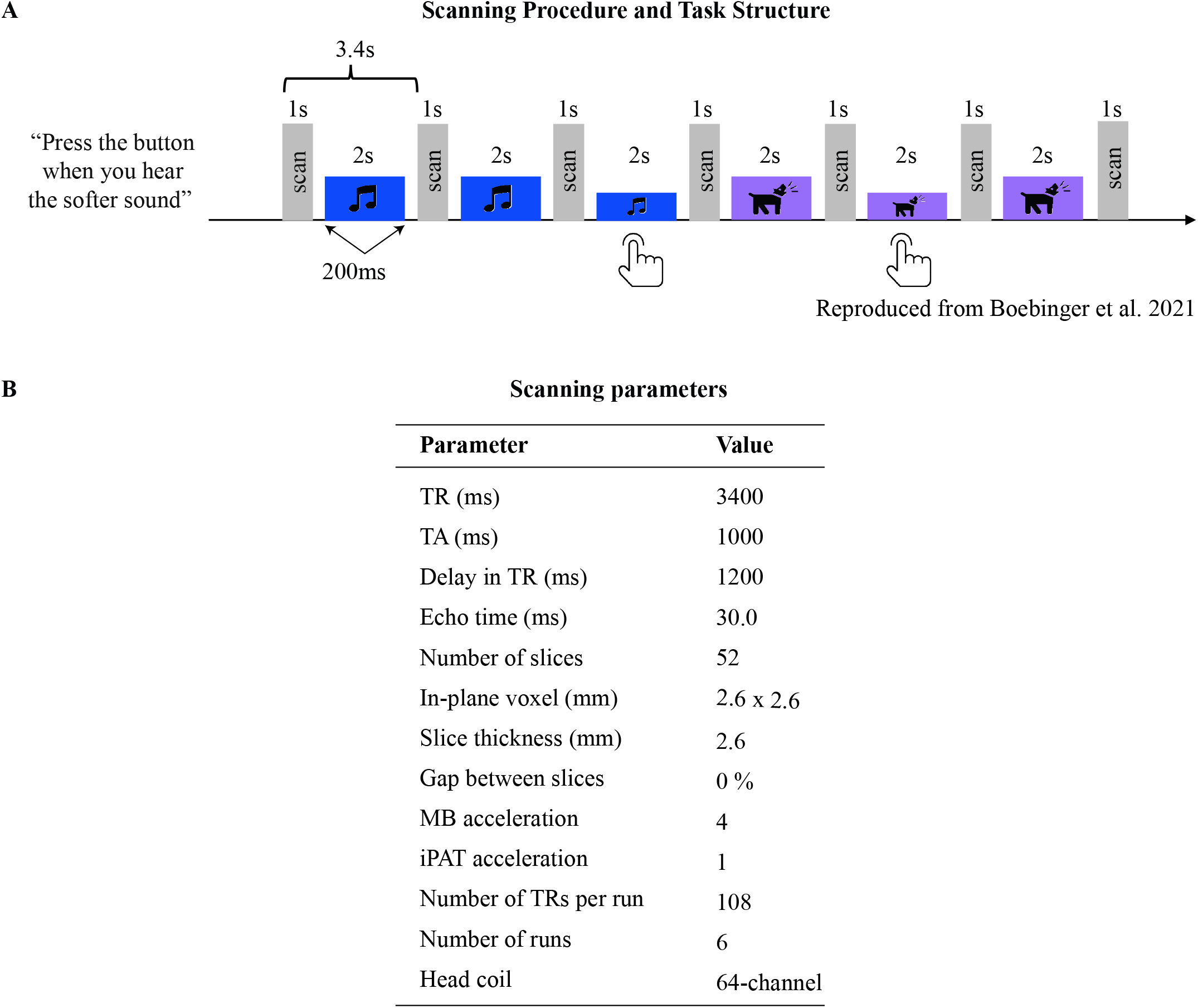
**A** Schematic of the scanning procedure. Sounds were presented between scans to mitigate the effects of scanner noise. Scans had an acquisition time (TA) of 1 second, and the 2-second sounds were padded by 200 milliseconds of silence at onset and offset, resulting in a total repetition time (TR) of 3.4 seconds. Each sound was played three times in succession, and either the second or third presentation was at a lower volume. Participants were tasked with pressing a button upon recognizing the quieter presentation. **B** fMRI scan parameters.

## Notes

https://labsites.rochester.edu/neural-prediction-decorrelation/

### Competing Interest Statement

The authors have declared no competing interest.

### Summary of Updates

Change from bar plots to box and whisker plots in Figure 4 D&E.

